# A new screening assay reveals that aminoglycoside antibiotics interfere with the Tau/MDM2 interaction

**DOI:** 10.1101/2024.10.24.619988

**Authors:** Martina Sola, Matteo Ciccaldo, Paolo Paganetti, Stéphanie Papin

**Author notes:** Corresponding author: Prof. Paolo Paganetti, Laboratories for Translational Research EOC, Room 102a, via Chiesa 5, CH-6500 Bellinzona, Switzerland, or.

## Abstract

*MAPT* gene mutations cause some neurodegenerative tauopathies and the *MAPT*-encoded protein Tau is deposited in neurofibrillary tangles, hallmarks of this disease family. In addition to its canonical function in regulating microtubule dynamics, Tau modulates chromatin compaction, gene expression, and the cellular response to DNA damage. During the DNA damage response, Tau positively modulates P53 by binding to MDM2 thereby preventing P53 inactivation and degradation. The aberrant presence of both MDM2 associated to neurofibrillary tangles and of P53 misfolding in brains affected by neurodegenerative diseases suggests that the sequestration of MDM2 may prevent P53 clearance in tauopathies and so contribute to progressive neuronal dysfunction and death. Following this evidence, a pharmacological inhibition of the Tau/MDM2 interaction may represent a viable strategy to reduce P53-dependent cell damage in brain disorders. With the screening FDA-approved drugs and natural compounds, we discovered that members of the aminoglycoside antibiotic family antagonize the Tau/MDM2 interaction. The use of these reagents may advance the understanding of the implication of the Tau/MDM2/P53 axis in neurodegeneration models. However, their unfavorable pharmacokinetic properties may limit their systemic use when targeting the brain.

## Introduction

Tau oligomerization and deposition in neurofibrillary tangles and neuropil threads is the hallmark of a group of neurodegenerative disorders termed tauopathies^1^. Tau deposits display disease-specific brain distribution and routes of propagation^2^. The most frequent tauopathy is Alzheimer’s Disease, where the extent of Tau deposits correlates with the clinical course. Notably, over forty autosomal dominant mutations in the *MAPT* gene encoding for Tau cause FTLD-Tau, a small group of progressive frontotemporal lobar degenerations^3,4^. The modification of Tau (phosphorylation or conformation) is assumed to impair the interaction of Tau with microtubules, to favor self-assembly into toxic Tau forms, and to initiate a cascade of gain-of-toxic events leading to neuronal dysfunction, clinical symptoms and premature mortality^5^.

In addition to its microtubule-stabilizing function, several additional roles of Tau have been reported. These include DNA binding^6^, DNA protection^6,7^, modulation of chromatin compaction^8–10^, and regulation of DNA damage response^11–14^. Moreover, in cells subjected to DNA damage, Tau modulates P53 stability and activity leading to a change in cell fate response^11^. Tau-dependent P53 modulation is achieved through the binding of Tau to MDM2^15^, which inhibits P53 ubiquitination *in vitro* in a manner sensitive to the presence of the P301L mutation linked to FTLD-Tau^11^. Notably, an abnormal accumulation of MDM2 in Alzheimer’s Tau tangles has been reported^15^, suggesting that MDM2 may be implicated in the disease course. These new functions of Tau also reinforce the possible implication of Tau in cancer and the implication of the Tau/MDM2/P53 axis in neurodegeneration.

A possible role of P53 in CNS disorders has been proposed^16^: whereas cancers are linked to P53 inactivation, in neurodegeneration the level and activity of P53 appear increased^17^. Genetic manipulation of P53 family members in mice affects aging and cognitive decline^18,19^. Loss of nuclear localization of P53 and P53 oligomers in the Alzheimer’s brain and brains of tauopathy mouse models is linked to a defective DNA damage response^12^, although a contribution of MDM2 in this process was not yet explored. Consistent with this, increased DNA damage is found in Alzheimer’s^20^ and persistent DNA damage causes neuronal senescence and upregulation of pro-inflammatory factors^21^. Finally, abnormal P53 species are considered Alzheimer’s biomarkers^22^. Importantly, nuclear Tau can form a complex with P53, Pin1 and PARN, a mRNA stability regulator, also for the P53 mRNA^23^. The cis/trans isomerase Pin1^24^ modifies Tau phosphorylation and is also sequestered into Tau lesions^25^ as is the case for MDM2. Moreover, Pin1 affects P53 phosphorylation, conformation, and the interaction with MDM2^26^. As a consequence, Tau and Pin1 facilitate P53-mediated PARN activation and the transcription deregulation in neurodegeneration and cancer^23^. P53 also associates to Tau oligomers^12^, suggesting that Tau intervenes on the P53-MDM2 axis by convergent mechanisms depending on the cell status.

To facilitate the identification of compounds interfering with the interaction between Tau and MDM2, we first developed an alphaLISA-based assay, which was then employed for the screening of natural compounds and FDA-approved drugs. This led to the discovery of inhibitors of the Tau/MDM2 complex enriched from Alzheimer’s brain homogenates. The hits, some identified in both libraries, are mainly members of the aminoglycoside antibiotic family.

## Materials and methods

### Preparation of human Alzhimer’s brain fractions

Frozen frontal cortex samples were obtained from completely anonymized donors through The Netherlands Brain Bank, Netherlands Institute for Neuroscience, Amsterdam (www.brainbank.nl). All donors signed a written informed consent for brain autopsy and the use of the material and clinical information for research purposes. Local guidelines foresee that work with completely anonymized human tissue samples, as is the case herein, does not require authorization from the local ethics committee. Brain tissue pooled from 4-5 Alzheimer’s donors was homogenized as described^37^ on ice in ∼9 volumes of filtered PHF buffer (10 mM Tris pH 7.4, 10% sucrose, 0.8 M NaCl and 0.1% sarkosyl, L9150-50G, Sigma) supplemented with protease and phosphatase inhibitor cocktails (S8820 & 04906845001, Sigma) in a glass Dounce homogenizer. Brain homogenates were briefly cleared by a centrifugation at 10’000 g for 10 min, 4°C. Sarkosyl was added to a 1% final concentration to the first supernatant (S0) and incubated for 90 min, room temperature, under agitation, before ultracentrifugation at 150’000 g for 75 min, 10°C. To remove sarkosyl, the P1 pellet was gently washed with cold PBS before repeating the ultracentrifugation. The P2 pellet was resuspended in 200 μl PBS supplemented with protease and phosphatase inhibitor cocktails and sonicated. The S2 supernatant was centrifuged at 10’000 g for 10 min, 4°C, and the Tau seed fraction S3 collected and stored frozen in aliquots. Total protein concentration was determined with the Pierce BCA protein assay kit (23227, ThermoFisher Scientific).

### Western Blot

Brain fractions were supplemented with SDS-PAGE sample buffer (1.5% SDS, 8.3% glycerol, 0.005% bromophenol blue, 1.6% β-mercaptoethanol and 62.5 mM Tris pH 6.8) and denatured for 10 min at 100 °C. For gel electrophoresis, 40 μg brain homogenate and 50 μg of all other fractions were loaded. After 10% SDS PAGE, PVDF membranes with transferred proteins were incubated with the anti-Tau primary antibodies: 0.2 μg/mL Tau13 (sc-21796 AF680, Santa Cruz) or 0.2 μg/mL AT8 (ab64193, Abcam). Primary antibodies were revealed with anti-mouse IgG coupled to IRDye 680RD (926–68070, Licor Biosciences) on a dual infrared imaging scanner (Odyssey CLx 9140, Licor Biosciences).

### MDM2 ELISA

Multititer 96 well plates (655101, Greiner) were coated with 7.4 ng/well S1 fraction or 16.7 ng/well S3 fraction in 50 mM carbonate buffer pH 9.6. Unspecific binding was blocked with 1% bovine serum albumin, 0.05% Tween-20 in PBS and washed were performed in 0.05% Tween-20 in PBS. Primary rabbit antibodies against MDM2 (D1V2Z, 86934, Cell Signaling; S395, PA5-13008, invitrogen) at 1:500 and secondary goat anti-rabbit antibody-HRP at 1:5000, were diluted in 0.1% bovine serum albumin, 0.05% Tween-20, PBS. The assay was developed with TMB (3,3’,5,5’-tetramethylbenzidine, 37574, Thermo Scientific) and stopped with 2 M phosphoric acid before reading the absorbance at 450 nm (Infinite 200Pro, Tecan).

### AlphaLISA

Tau protein was determined by alphaLISA (Tau-AL271C, PerkinElmer) following the instructions of the manufacturer. A home-made assay was developed for the Tau/MDM2 complex as described^15^, using D1V2Z-CaptS (anti-MDM2 D1V2Z coupled to CaptSure following the instructions of the manufacturer, 6300007, Expedeon) and CaptSure-conjugated acceptor beads (ALSU-ACAB, PerkinElmer), paired to biotinylated anti-Tau HT7 (MN1000B, ThermoFisher) and streptavidin-conjugated donor beads (6760002, PerkinElmer). The assays were performed in 384-well plates (6005350, PerkinElmer). Ten μL samples and 5 μL of a mix containing biotinylated antibody (1 nM final concentration), antibody-CaptS bound to acceptor beads (10 μg/mL final concentration) were incubated in HiBlock for 1 h at room temperature in the dark. This was followed by the addition of 5 μL of donor beads (20 μg/mL final concentration) in HiBlock for another 1 h incubation. Measurement was done with a multi-plate reader (Victor Nivo, PerkinElmer) and data were analyzed using the provided software (Victor Nivo, PerkinElmer) with excitation time of 50 msec and 575/110 nm emission time of 700 msec.

### Compound screening

Compounds were obtained from two commercial libraries. First, the screen-well FDA approved drug library (FDA BML-2843, V1.4 Dec 19, 2019, Enzo Life Sciences) containing ∼770 compounds in eleven 96-well plates at 10 mM in DMSO and ∼30 compounds at 10 mM in water). Second, the screen-well natural product library (NP BML-2865, V7.6 Nov 18, 2020, Enzo Life Sciences) containing ∼500 compounds in six 96-well plates at 2 mg/mL in DMSO (4 compounds in water). Compounds were tested in the primary screen at 50-100 μM in monoplicate. Each of the 17 multititer plates screened contained 3 wells without the analyte S1 (the mean value of which was defined as 0%) and 6-12 wells with S1 diluted 1:450 to 13.1 μg/mL and DMSO only as the main vehicle (the mean value of which was defined as 100%). Positive primary hits reduced the Tau/MDM2 alphaLISA signal in the range 50-105%. The hits were retested under the same conditions utilizing the Tau/MDM2 alphaLISA and the Tau alphaLISA as counter-screen. Confirmed hits reduced the amount of Tau/MDM2 complex detected in the range 50-105% and the Tau alphaLISA signal <25%. Final confirmation was performed in triplicate for selected hit compounds.

### Statistics and Reproducibility

Statistical analysis was performed with GraphPad Prism version 10.3 using the method specified in the legend of each figure.

## Results

### An assay to detect the Tau/MDM2 complex in brain homogenates

We reported the biochemical detection of the Tau/MDM2 complex in a human brain homogenate fraction utilizing a highly sensitive alphaLISA assay assembled with antibodies against each one of the two proteins^15^. The Tau/MDM2 assay was built with alphaLISA acceptor beads coated with the anti-MDM2 D1V2Z antibody in combination with alphaLISA streptavidin-donor beads and biotinylated anti-Tau Tau13 antibody. Close-proximity between donor and acceptor beads, due to the presence of the Tau/MDM2 complex, results in oxygen radical transfer from the laser-excited donor beads to the acceptor beads that will emit fluorescence. To obtain the analyte (Tau/MDM2 complex), we took advantage of a sequential centrifugation method to enrich for endogenous Tau oligomers from the Alzheimer’s brain (**Fig 1A**). We saved the intermediate fractions and tested them for the presence of Tau by western blot (**Fig 1B**), and of MDM2 by direct immune detection of human brain proteins coated to multititer plates (**Fig 1C**). The soluble S1 (5.9 mg/mL) and S3 (4.2 mg/mL) fractions contained detectable amounts of Tau and MDM2. So, we tested them for the presence of the endogenous Tau/MDM2 with the Tau/MDM2 assay. Three dilutions of the S1 and S3 fractions were tested against the signal of negative controls obtained by omitting one or the other detection antibodies in the assay (**Fig 1D**). Based on these data, we selected the highest signal to noise condition, obtained from the S1 fraction diluted 1:125 (47.5 μg/mL), for the compound screening. It should be noted that for the negative controls, a larger unspecific signal was obtained in the absence of the D1V2Z antibody when compared to the absence of the Tau13 antibody. This observation, consistent with others done in the laboratory, suggests non-specific binding of Tau to solid-phase surfaces, also under stringent conditions.

**Figure 1.**
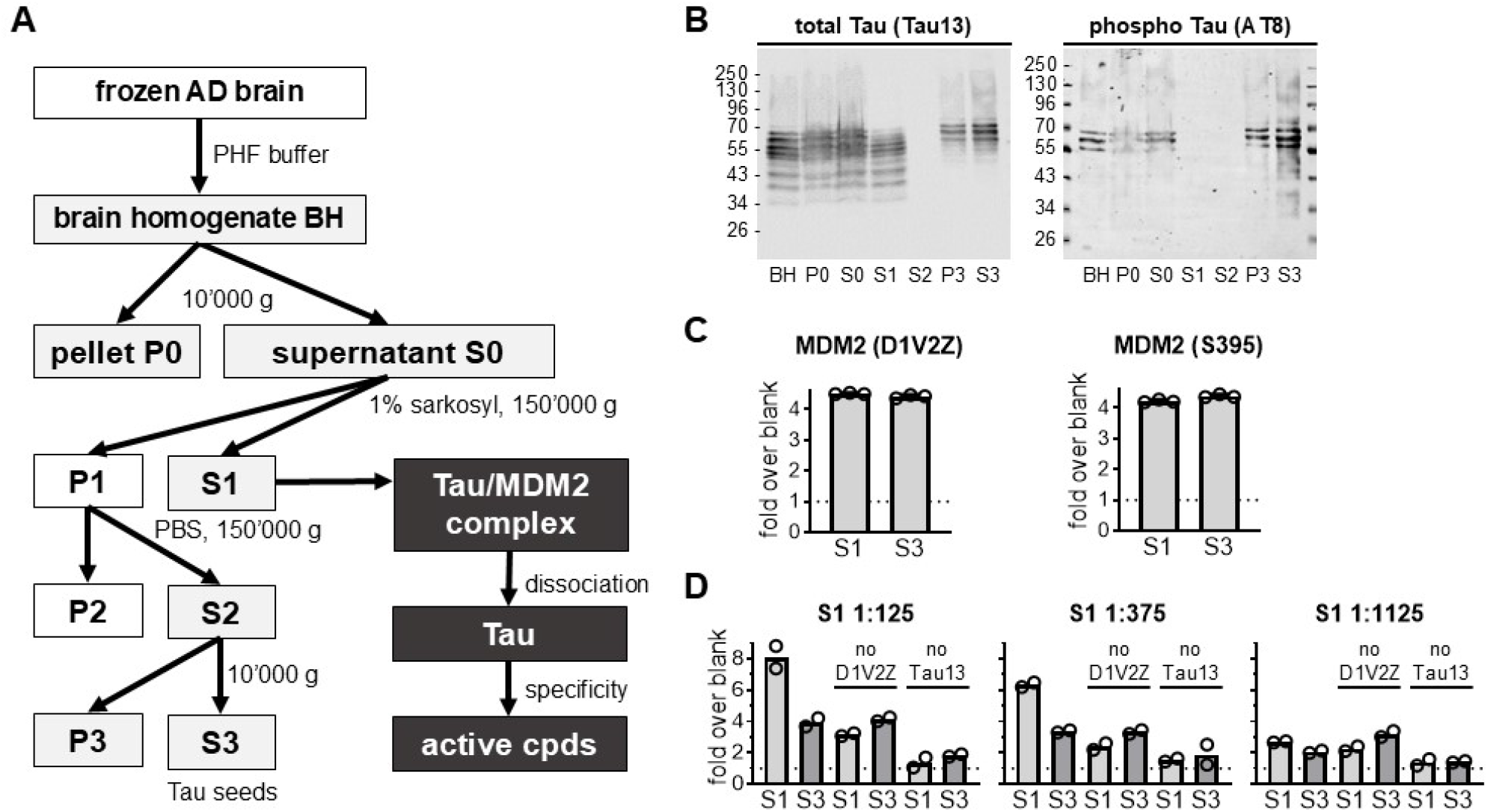
Assay for endogenous Tau/MDM2 complex from human brain. **A**. Scheme of the Tau seed procedure used for enriching the Tau/MDM2 complex from frozen Alzheimer’s brain tissue and procedure to screen and counter-screen test compounds in the S1 150’000 g supernatant fraction. **B**. The indicated human brain fractions were analyzed by western blot with the Tau13 antibody against total Tau or the AT8 antibody against phosphoTau. Primary antibodies were revealed with anti-mouse-IRDye700CW secondary antibody. **C**. Detection of MDM2 in the S1 and S3 fractions by ELISA with the MDM2 rabbit antibodies D1V2Z or S395. Values are given as fold over empty wells (blank), mean ± SD of n=3 wells. **D**. Detection of the endogenous Tau/MDM2 complex in the S1 and S3 human brain fractions diluted as indicated by alphaLISA. If indicated either the primary antibody D1V2Z or Tau13 were omitted from the reaction as negative control for unspecific signals. Values are given as fold over buffer alone (blank), mean of n=2 samples.

### Compound screening

The Tau/MDM2 alphaLISA was first tested for the tolerability towards the solvent used to prepare most of the stock solutions, when not dissolved in water. For this, the Tau/MDM2 complex present in the S1 fraction was determined in the presence of 0.1%, 0.3% and 1% DMSO and compared to its absence. Highly significant signal to noise ratios were found for all conditions, whereby a small, but significant, dose-dependent increase in this ratio was observed for the two higher DMSO concentrations (**Fig 2A**). We concluded that the presence ≤ 1% DMSO in the reaction solution did not compromise the quality of the assay; this allowed for testing compounds diluted not less than 100-times starting from their DMSO stock solutions. The reproducibility of the assay over a 96-well multititer plate was tested in the presence of 1% DMSO by analyzing 48 wells each in the presence (positive signal) or absence (negative signal) of the S1 homogenate. We determined a Z’ value of 0.83 i.e., in a range suited for screening small molecular weight compounds (**Fig 2B**). The primary screen was run on two commercial libraries provided by Enzo: the screen-well FDA approved drug library V1.4 (∼800 compounds, 10 mM in DMSO) and the screen-well natural product library V7.6 (∼500 compounds, 2 mg/mL in DMSO). The 1:125 diluted S1 fraction was incubated for 30 min in the presence of compounds diluted at 50-100 μM in monoplicate before determination of the Tau/MDM2 complex. The screening of the ∼1300 compounds distributed in 17 microtiter plates was performed manually (see an example of a microtiter plate in **Fig 2C**). We defined 0% as the mean of background signals obtained from wells devoid of the analyte (S1 fraction) and 100% as the mean signal determined from wells containing S1 and 1% DMSO. These controls were present in each screening plate. The great majority of signals generated when testing the compounds distributed in the range 100 ± 50% (**Fig 2D**). We identified 95 hit compounds that reduced the Tau/MDM2 signal by >50% (but not >105%) of that determined in the DMSO only controls (**Fig 2D**). Notably, among the 17 plates composing the two libraries, 35 wells did not contain compounds and, as expected for vehicle-only conditions, none of these wells were found positive. The activity of the hit compounds was confirmed in triplicates with the Tau/MDM2 alphaLISA and counter-screened with a Tau-specific alphaLISA. To exclude possible interference with the alphaLISA technology, compounds reducing the Tau signal by >25% were excluded from further analysis (**Fig 2E**). One of the hits, gentamycin sulfate, was purchased from a different provider and tested at different concentrations. Gentamycin was found to reduce the Tau/MDM2 signal in a dose-dependent manner at concentrations >10 nmol/L (**Fig 2F**).

**Figure 2.**
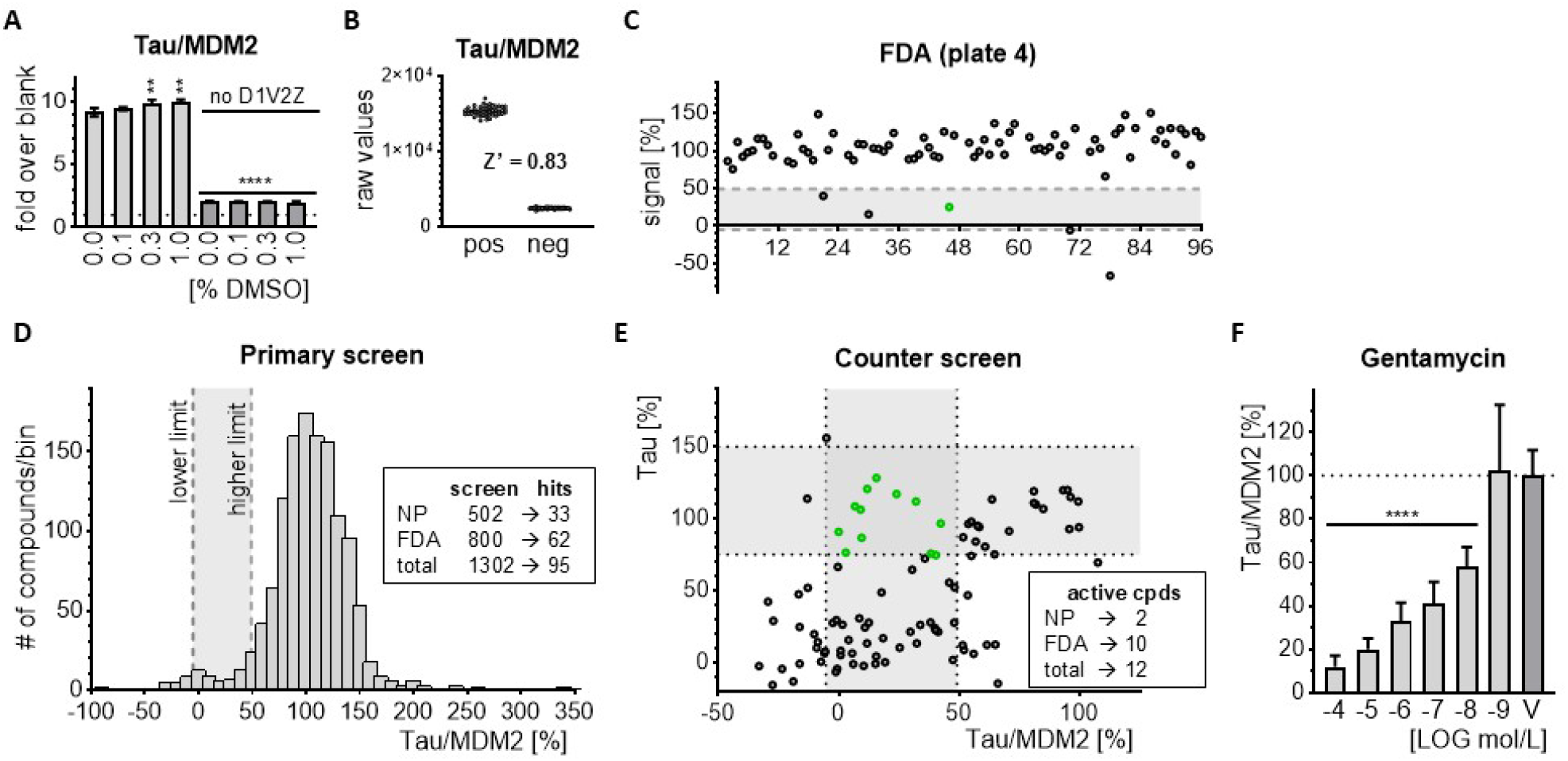
Screening for compounds dissociating endogenous Tau/MDM2. **A**. The amount of Tau/MDM2 complex in the S1 brain fraction was tested by alphaLISA in the presence of different amounts of DMSO as indicated. Unspecific signals were obtained by omitting the D1V2Z antibody as indicated. Values are given as fold over empty wells (blank), mean ± SD of n=3 wells. Ordinary one-way ANOVA (p <.0001) and Šídak’s multiple comparisons test comparing no DMSO condition to all other conditions; ** p <.01, **** p <.0001. **B**. Tau/MDM2 raw values obtained from 48 wells each in presence (pos) or absence (neg) of the S1 brain fractions. The two conditions were distributed randomly across a multititer plate. Z’ was calculated with the formula 1-[(3xSDpos + 3xSDneg)/(mean pos – mean neg)]. **C**. Example of a screening plate (FDA drug plate #4). Values are reported as percent (0% is no S1, 100% is S1 no compound). Dashed lines show the 50-105% range utilized for scoring positive primary hits. In green the value for gentamycin sulfate in well D10. **D**. Distribution of the activity for the ∼1300 compounds screened. Shown is the number of compounds contained in the indicated 10%-wide bins. Dashed lines show the 50-105% range utilized for scoring positive primary hits. **E**. Secondary screen (Tau/MDM2 alphaLISA) and counter-screen (Tau alphaLISA) for 95 primary hits. Shown are the mean percent values obtained for each compound in both assays. Vertical dashed lines (50-105% Tau/MDM2 value) and horizontal dashed lines (75-150% Tau value) define the region including the 12 confirmed hits reducing Tau/MDM2 but not Tau. **F**. Dose-dependent reduction of Tau/MDM2 complex in the presence of decreasing amounts of gentamycin sulfate. Values are given mean ± SD of n=6-9 wells (n=2 for 10^-9^ mol/L). Ordinary one-way ANOVA (p <.0001) and Šídak’s multiple comparisons test comparing vehicle (V) to all other conditions; **** p <.0001.

In summary, we identified 10 compounds, which reduced the amount of endogenous Tau/MDM2, but not of Tau, in human brain homogenates (**Fig 3A**). Among the ∼500 natural products tested, we found only two active compounds: neomycin (well NP 1-H6) and kanamycin (well NP 2-H10). The same two compounds were also identified among the ten active compounds of the ∼800 FDA drug library (wells 2-F11 and 8-F02, respectively). Capreomycin in well 6-F7 of the FDA library reduced the Tau/MDM2 signal by 93%, whereas the same compound in well NP 4-G8 of the natural product library reduced the Tau/MDM2 signal by 30% and resulted negative. In total, the two libraires had ∼40 duplicate compounds. So, whereas 100% of the natural product hits were also FDA hits, only 8% of the natural compounds were present in the FDA drug collection, validating the quality of the screen. Moreover, among the ten hits with distinct chemical structures, five of them were aminoglycoside antibiotics (**Fig 3B**).

**Figure 3.**
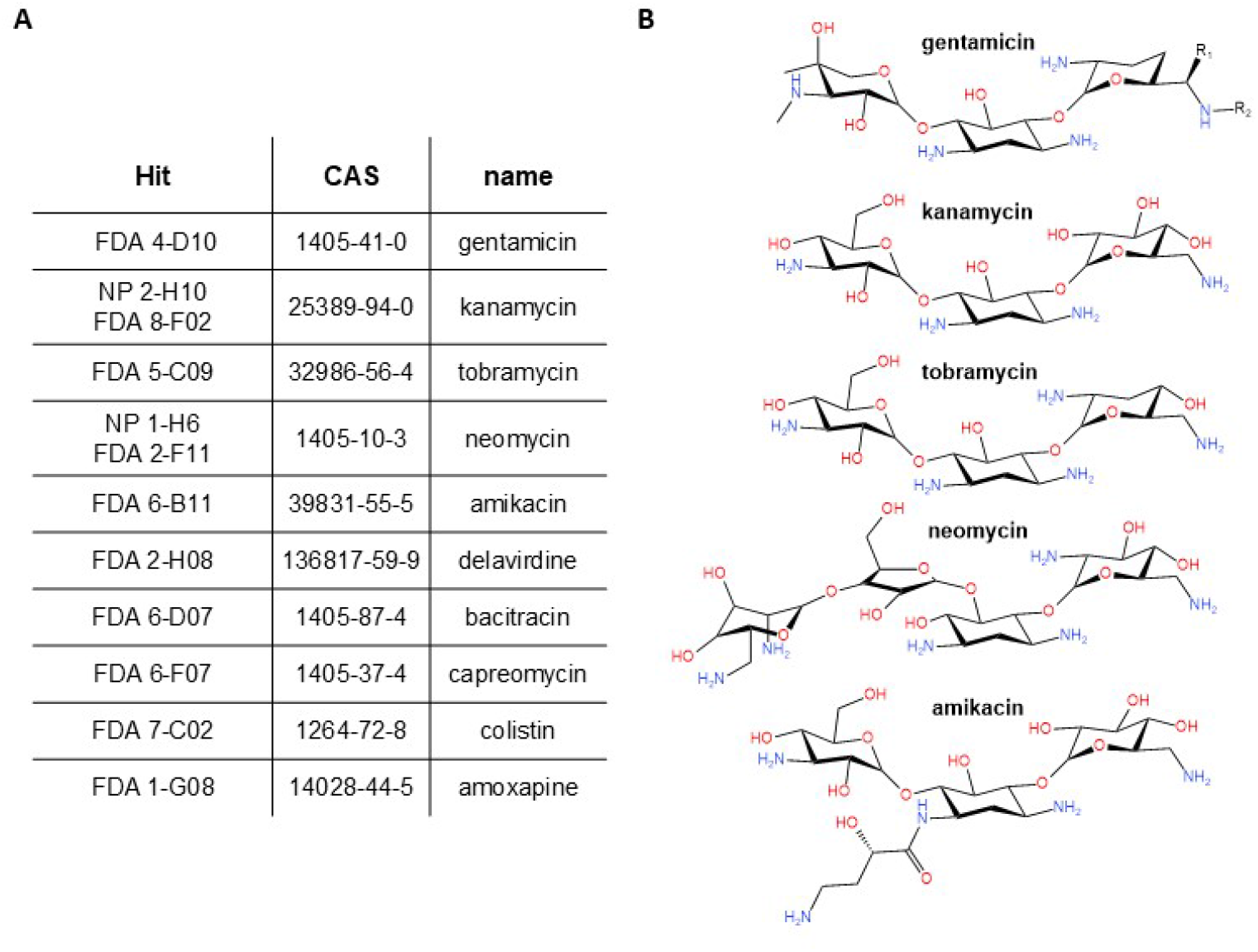
Confirmed hit compounds reducing the Tau/MDM2 complex. **A**. Shown are the screening plate location of the ten specific antagonists of the endogenous human brain Tau/MDM2 complex, their unique Chemical Abstracts Service (CAS) registry number and their trade name. **B**. Chemical structures of the five aminoglycoside antibiotics found among the 10 confirmed hits.

## Discussion

We previously reported by cross-validation with orthogonal technologies that Tau binds to the E3 ubiquitin-ligase MDM2 *in vitro*, in cultured human cells and in human brain tissue^15^. The MDM2 interactome includes 200 proteins such as ribosomal proteins, transcription factors, tumor suppressors, DNA repair mediators, cell fate modulators, E2 ubiquitin-protein ligases and other regulators of cell function^27^. The large number of binding partners may result from the scaffold organization of MDM2, encompassing three structured domains spaced by disordered polypeptides. This lack of binding specificity contrasts with the selective and tightly regulated functions attributed to MDM2. Perhaps, cell-specific post-translational modifications and conformations may reduce the MDM2 interacting partners^28^. Also, absence of binding partners in differentiated cells may limit the spectrum of MDM2 interactions and activities. We now extend our results with the identification of small MW compounds, in particular aminoglycoside antibiotics, which present the ability to dissociate *in vitro* the endogenous Tau/MDM2 complex detected in human Alzheimer’s brain homogenates.

The central acidic domain of MDM2 plays a crucial part in the interaction with Tau^15^ as well as with numerous other proteins^27^. The acidic domain drives also MDM2 homodimerization^29^ and its intramolecular interaction with the RING domain impairs binding to the proteasome^30^. In cooperation with the canonical N-terminal P53 binding domain, the central domain of MDM2 induces a change in the structure of P53 in a CKIδ phosphorylation-dependent manner^30–32^. It is thus expected that Tau may regulate many MDM2 functions in addition to P53 ubiquitination^15^ and stability^11^. In fact, many proteins bind the central region of MDM2 and regulate its activity. A prime example is the tumor suppressor ARF, which targets MDM2 to the nucleolus and inhibits its E3 ligase function^33,34^. The herein reported identification of compounds interfering with the binding of Tau to MDM2 may become instrumental in analyzing which MDM2 functions may be regulated by Tau. We cannot exclude at this point that these compounds may interfere with the binding of other MDM2 partners.

However, we should also consider how Tau may bind MDM2 at the molecular level. Tau binds to MDM2 through its microtubule-binding domain (MBD)^15^. For the naturally unfolded protein Tau^5^, the MBD acquires a more defined structure when binding to microtubules^35^. Tau binding to microtubules relies on electrostatic forces between the net positive charge of the MBD and the rather acidic character of the microtubule surface^37^. In fact, phosphorylation of the MBD inhibits its binding to microtubules and may eventually lead to pathological Tau self-assembly^5^, also supported by the fact that most autosomal dominant FTLD-Tau mutations target the MBD^36^. Notably, the many functional groups of aminoglycoside antibiotics may orient themselves in opposing directions (**Fig. 3B**) causing a net distribution of positive and negative charges possibly fitting between the negatively charged acidic domain of MDM2 and the positively charged MBD of Tau.

The use of Tau/MDM2 complex inhibitors may facilitate our understanding of the role of the Tau/MDM2/P53 axis in neurodegenerative processes. However, their unfavorable pharmacokinetic properties may limit their systemic use when targeting the brain.

## Acknowledgments

We thank the whole laboratory for support and advice during this study. This study was supported by generous grants from the Novartis Foundation and the Gelu Foundation.

## Author contribution

Conceptualization: SP, PP

Methodology: MS, MC

Investigation: MS, MC

Supervision: SP, PP

Writing (original draft and figures): PP

Writing (review & editing): MS, MC, PP, SP

## Data and materials availability

all data generated or analyzed during this study are included in this published article.

